# Host phenology can drive the evolution of intermediate virulence strategies in some obligate-killer parasites

**DOI:** 10.1101/2021.03.13.435259

**Authors:** Hannelore MacDonald, Erol Akçay, Dustin Brisson

**Author notes:** **Cite as:** MacDonald H, Akçay E, Brisson D (2021) Host phenology can drive the evolution of intermediate virulence strategies in some obligate-killer parasites. bioRxiv, 2021.03.13.435259, ver. 8 peer-reviewed and recommended by Peer Community in Evolutionary Biology. https://doi.org/10.1101/2021.03.13.435259. This article has been peer-reviewed and recommended by Peer Community In Evolutionary Biology (https://doi.org/10.24072/pci.evolbiol.100129).

## Abstract

The traditional mechanistic trade-offs resulting in a negative correlation between transmission and virulence are the foundation of nearly all current theory on the evolution of parasite virulence. Several ecological factors have been shown to modulate the optimal virulence strategies predicted from mechanistic trade-off models, but these ecological factors have not yet been shown to be sufficient to explain the intermediate virulence strategies observed in any natural system. The timing of seasonal activity, or phenology, is a common factor that influences the types and impact of many ecological interactions but is difficult to incorporate into virulence evolution studies. We develop a mathematical model of a disease system with seasonal host activity to study the evolutionary consequences of host phenology on the virulence of obligate-killer parasite. Results from this model demonstrated that seasonal host activity is sufficient to drive the evolution of intermediate parasite virulence in some types of natural disease systems, even when a traditional mechanistic trade-off between transmission and virulence is not assumed in the modeling framework. The optimal virulence strategy in these systems can be determined by both the duration of the host activity period as well as the variation in the host emergence timing. Parasites with low virulence strategies are favored in environments with long host activity periods and in environments in which hosts emerge synchronously. The results demonstrate that host phenology can be sufficient to select for intermediate optimal virulence strategies, providing an alternative mechanism to account for virulence evolution in some natural systems.

## Introduction

The evolutionary causes and consequences of parasite virulence remain enigmatic despite decades of research. It was once thought that parasites continue to evolve ever lower levels of virulence to preserve their primary resource for future parasite generations (Smith, 1904). However, the idea that parasites would limit damaging their host for the benefit of future generations violates multiple core principles of our modern understanding of evolutionary biology (Hamilton, 1964a,b). Natural selection, as framed in the modern evolutionary synthesis, favors traits that improve short-term evolutionary fitness even if those traits negatively impact the environment for future generations (Hamilton, 1964a,b). Thus, the level of virulence that maximizes parasite fitness is favored by natural selection despite its impact on the host population (RM Anderson and May, 1982). Identifying environmental conditions and mechanistic constraints that drive the evolution of the vast diversity of parasite virulence strategies observed in nature has been an important research focus for decades.

Establishing that mechanistic trade-offs constrain parasite virulence strategies was a key breakthrough that propels virulence evolution research to this day. In the now classic paper, Anderson and May demonstrated that within-host parasite densities increase the probability of transmission to a new host - a component of parasite fitness - but also shorten infection duration resulting in fewer opportunities for transmission to naive hosts (RM Anderson and May, 1982). That is, parasites can only produce more infectious progeny if they cause host damage by utilizing more host resources. Mechanistic trade-offs results in a negative correlation between virulence and transmission and remain the foundational theoretical framework used to account for the evolution of parasite virulence strategies observed in nature (Alizon et al., 2009; Cressler et al., 2016).

Many ecological factors and environmental conditions have been shown to alter the optimal virulence strategies driven by mechanistic trade-offs within models. For example, it is well-established that varying environmental conditions, such as the extrinsic host death rate, often shift the optimal virulence strategy governed by a mechanistic trade-off (RM Anderson and May, 1982; Cooper et al., 2002; Gandon et al., 2001; Lenski and May, 1994). However, no environmental condition, *in the absence* of an explicitly modeled negative correlation between virulence and transmission, has been shown to select for intermediate virulence.

The timing of seasonal activity, or phenology, is an environmental condition affecting all aspects of life cycles, including reproduction, migration, and diapause, in most species (J Anderson et al., 2012; Elzinga and e ae, 2007; Forrest and Miller-Rushing, 2010; Lustenhouwer et al., 2018; Novy et al., 2013; Park, 2019; Pau et al., 2011). The phenology of host species also impacts the timing and prevalence of transmission opportunities for parasites which could alter optimal virulence strategies (Altizer et al., 2006; Biere and Honders, 1996; Gethings et al., 2015; Hamer et al., 2012; MacDonald et al., 2020; Martinez, 2018; McDevitt-Galles et al., 2020; Ogden et al., 2018). For example, host phenological patterns that extend the time between infection and transmission are expected to select for lower virulence, as observed in some malaria parasites (*Plasmodium vivax*). In this system, high virulence strains persist in regions where mosquitoes are present year-round while low virulence strains are more common in regions where mosquitoes are nearly absent during the dry season(MT White et al., 2016). While host phenology likely impacts virulence evolution in parasites (Berg et al., 2011; Donnelly et al., 2013; King et al., 2009; Sorrell et al., 2009), it remains unclear whether this environmental condition can have a sufficiently large impact to select for an intermediate virulence phenotype in the absence of a mechanistic trade-off.

Here we investigate the impact of host phenology on the virulence evolution of an obligate-killer parasite. We demonstrate that intermediate virulence is adaptive when host activity patterns are highly seasonal, establishing that environmental context alone is sufficient to drive the evolution of intermediate virulence in disease systems that conform to the assumptions of the model. Further, multiple features of host seasonal activity, including season length and the synchronicity at which hosts first become active during the season, impact the optimal virulence level of parasites. These results provide an alternative framework that can account for virulence evolution in some natural systems.

### Model description

The model describes the transmission dynamics of a free-living, obligate-killer parasite that infects a seasonally available host (Figure 1). The host cohort, 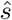, enters the system at the beginning of the season over a period given by the function *g*(*t, t*_*l*_). Hosts, *s*, have non-overlapping generations and are alive for one season. The parasite, *v*, infects hosts while they are briefly susceptible early in their development (*e*.*g*. baculoviruses of forest *Lepidoptera* (Baltensweiler et al., 1977; Bilimoria, 1991; Dwyer, 1994; Dwyer and JS Elkinton, 1993; Woods and J Elkinton, 1987) and univoltine insects parasitized by ichneumonids (Campbell, 1975; Delucchi, 1982; Kenis and Hilszczanski, 2007)). The parasite must kill the host to release new infectious progeny. The parasite completes one round of infection per season because the incubation period of the parasite is longer than the duration of time the host spends in the susceptible developmental stage. This transmission scenario occurs in nature if all susceptible host stages emerge over a short period of time each season so that there are no susceptible host stages available when the parasite eventually kills its host. Parasites may also effectively complete only one round of infection per season if the second generation of parasites do not have enough time in the season to complete their life cycle in the short-lived host.

**Figure 1.**
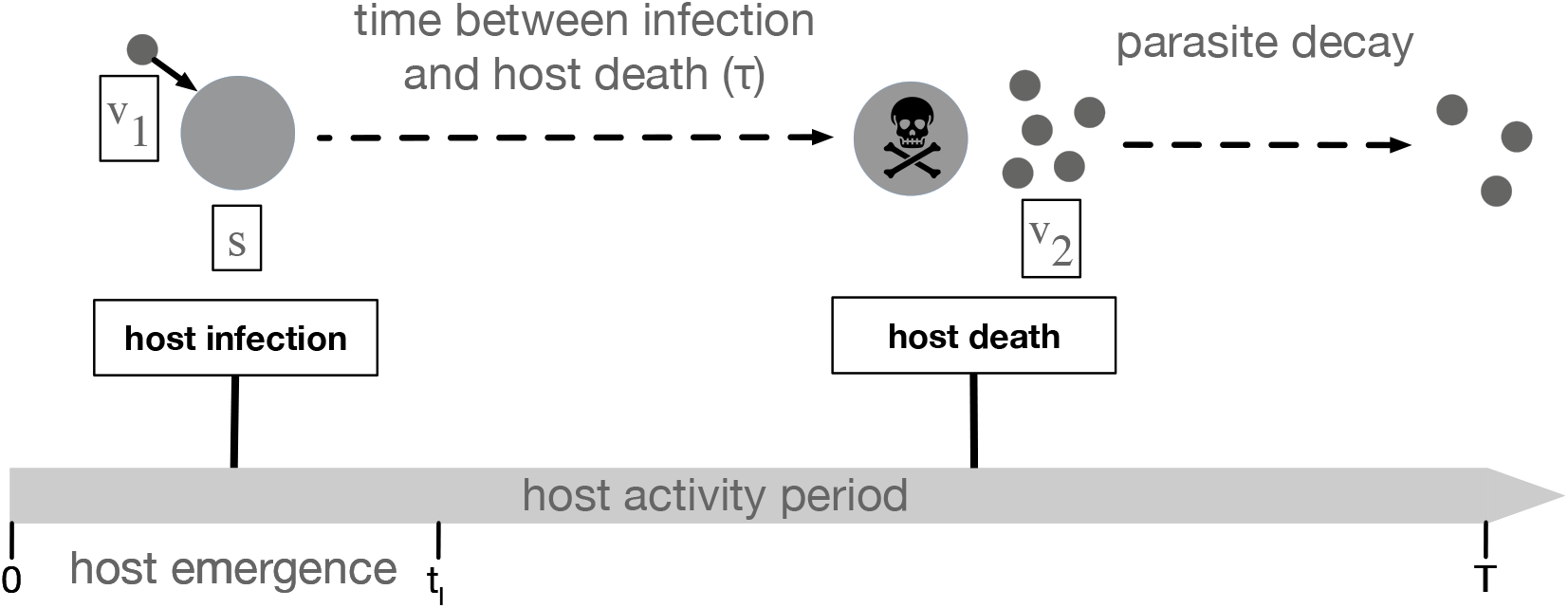
Infection diagram. The host cohort, 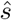, emerges from time *t* = 0 to *t* = *t*_*l*_, all *v*_1_ parasites emerge at *t* = 0. Hosts do not reproduce during the season. Infections generally occur early in the season when host density is high. Parasite-induced host death occurs after time *τ*, at which point new parasites, *v*_2_ are released. *v*_2_ decays in the environment from exposure. Parasites only have time to complete one round of infection per season. *v*_2_ parasites in the environment at *t* = *T* will carryover and emerge at the beginning of the next season.

We ignore the progression of the susceptible stage, *s*, to later life stages as it does not impact transmission dynamics. To keep track of these dynamics, we refer to the generation of parasites that infect hosts in the beginning of the season as *v*_1_ and the generation of parasites released from infected hosts upon parasite-induced death as *v*_2_. *τ* is the delay between host infection by *v*_1_ and host death when *v*_2_ are released. *τ* is equivalent to virulence where low virulence parasites have long *τ* and high virulence parasites have short *τ*. The initial conditions in the beginning of the season are *s*(0) = 0, *v*_1_(0^+^) = *v*_2_(0^*−*^), *v*_2_(*τ*) = 0. The transmission dynamics in season *n* are given by the following system of delay differential equations (all parameters are described in Table 1):

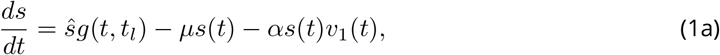

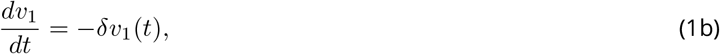

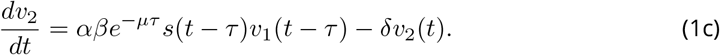

where *µ* is the host death rate, *δ* is the decay rate of parasites in the environment, *α* is the transmission rate, *β* is the number of parasites produced upon host death. We make the common assumption for free-living parasites that the removal of parasites through transmission (*α*) is negligible (RM Anderson and May, 1981; Caraco and Wang, 2008; Dwyer, 1994), *i*.*e*. (1b) ignores the term *−αs*(*t*)*v*_1_(*t*).

**Table 1.** Model parameters and their respective values.

| Parameter | Description | Value |
| --- | --- | --- |
| $s$ | susceptible hosts | state variable |
| $v_1$ | parasites that infect hosts in current season | state variable |
| $v_2$ | parasites released in current season | state variable |
| $t_l$ | length of host emergence period | time (varies) |
| $T$ | season length | time (varies) |
| $\hat{s}$ | emerging host cohort size | $10^8$ hosts |
| $\alpha$ | transmission rate | $10^{-8}/(\text{parasite} \times \text{time})$ |
| $\beta$ | number of parasites produced upon host death | parasites (varies) |
| $\delta$ | parasite decay rate in the environment | 2 parasites/parasite/time |
| $\mu$ | host death rate | 0.5 hosts/host/time |
| $\tau$ | time between host infection and host death (1/virulence) | time (evolves) |

The function *g*(*t, t*_*l*_) captures the per-capita host emergence rate by specifying the timing and length of host emergence. We use a uniform distribution (*U* (*•*)) for analytical tractability, but other distributions can be used.

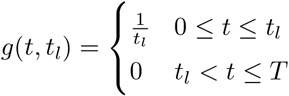

*t*_*l*_ denotes the length of the host emergence period and *T* denotes the season length. The season begins (*t*_0_ = 0) with the emergence of the susceptible host cohort, 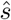. The host cohort emerges from 0 ≤ *t* ≤ *t*_*l*_. *v*_2_ parasites remaining in the system at *t* = *T* give rise to next season*’*s initial parasite population 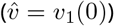. Parasites that have not killed their host by the end of the season do not release progeny. Background mortality arises from predation or some other natural cause. We assume that infected hosts that die from background mortality do not release parasites because the parasites are either consumed or the latency period corresponds to the time necessary to develop viable progeny (Wang, 2006; NJ White, 2011). We ignore the impact of infection for host demography and assume 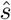 is constant each year (*e*.*g*. a system where host regulation by parasites is negligible). We solve equations 1a-c analytically Appendix A.

### 0.0.1 Parasite fitness

A parasite introduced into a naive host population persists or goes extinct depending on the length of the host emergence period and season length. The stability of the parasite-free equilibrium is determined by the production of *v*_2_ resulting from infection of *s* given by

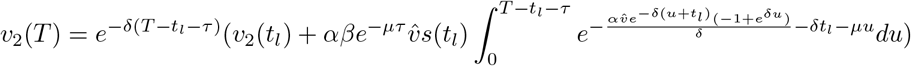

when *τ < T − t*_*l*_ and by

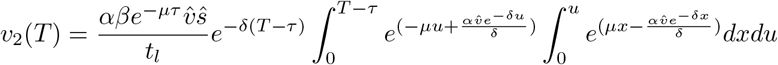

when *τ > T − t*_*l*_.

The parasite-free equilibrium is unstable and a single parasite introduced into the system at the beginning of the season will persist if the density of *v*_2_ produced by time *T* is greater than or equal to 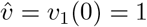 (*i*.*e. v*_2_(*T*) ≥ 1, modulus is greater than unity). This expression is a measure of a parasite*’*s fitness when rare given different host phenological patterns. See Appendix A for details of the analytical solution.

### 0.0.2 Parasite evolution

To study how parasite traits adapt given different seasonal host activity patterns, we use evolutionary invasion analysis (Geritz et al., 1998; Metz et al., 1992). We first extend system (1) to follow the invasion dynamics a rare mutant parasite

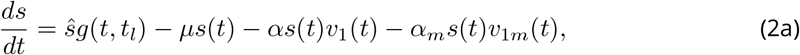

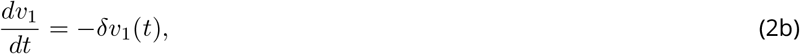

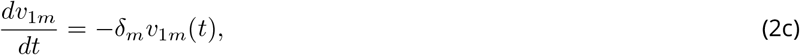

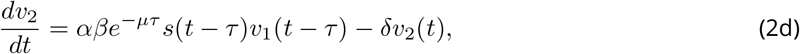

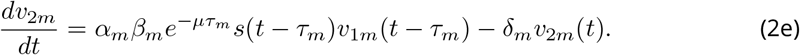

where *m* subscripts refer to the invading mutant parasite and its corresponding traits. See Appendix B for details of the time-dependent solutions for equations (2a-2e).

The invasion fitness of a rare mutant parasite depends on the density of *v*_2*m*_ produced by the end of the season (*v*_2*m*_(*T*)) in the environment set by the resident parasite at equilibrium density 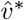. The mutant parasite invades in a given host phenological scenario if the density of *v*_2*m*_ produced by time *T* is greater than or equal to the initial *v*_1*m*_(0) = 1 introduced at the start of the season (*v*_2*m*_(*T*) ≥ 1). When *τ < T − t*_*l*_, mutant invasion fitness can be found using

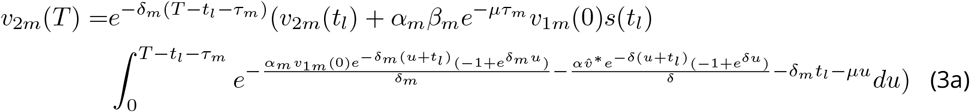

When *τ > T − t*_*l*_, mutant invasion fitness can be found using

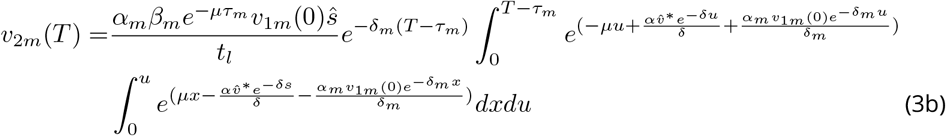

To study the evolution of virulence traits, we first assume all other resident and mutant traits are identical (*e*.*g. α* = *α*_*m*_). Note that when there is no trade-off between *β* and *τ*, the parasite growth rate in the host is essentially the trait under selection. That is, *β* is constant regardless of *τ*, thus the trait that is effectively evolving is the rate that new parasites are assembled in between infection and host death (*e*.*g*. long *τ* corresponds to slow assembly of new parasites.) To find optimal virulence for a given host phenological scenario, we find the uninvadable trait value that maximizes (3). That is, the virulence trait, *τ* ^∗^, that satisfies

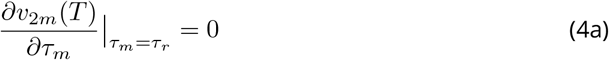

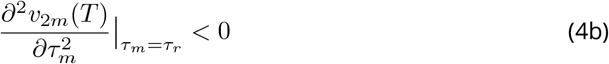

Note that the measure in equation (3) incorporates the effect of the resident on the population state (number of susceptibles over one season), which means that it is not a measure of *R*_0_ (which by definition assumes a non-disease environment). Thus, we can use *v*_2*m*_(*T*) as defined in (3) as a maximand in evolutionary dynamics (Lion and Metz, 2018).

To study the impact of mechanistic trade-offs between transmission and virulence on virulence evolution, we assume that the number of parasites produced at host death is a function of the time between infection and host death (*β*(*τ*)). For example, mutant invasion fitness for *τ < T − t*_*l*_ can be found using

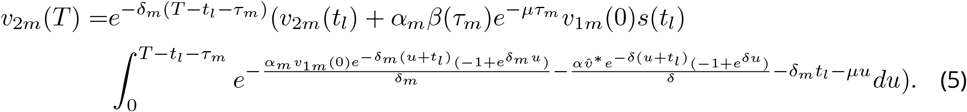

We then find *τ* ^∗^ that satisfies (4a) and (4b) using equation (5).

## Results

Host phenology is sufficient to drive the evolution of intermediate virulence in systems that conform to the assumptions of the model. Host phenology is composed of the duration of the activity period and the distribution of initial emergence times, both of which impact the optimal parasite virulence level. Temporally constrained host activity periods within each season can select against both extremely high and extremely low virulence levels resulting in an intermediate optimal level of virulence. Low virulence is selected against as parasites that do not kill the infected host prior to the end of the host activity period fail to produce progeny and thus have no evolutionary fitness. By contrast, highly-virulent parasites kill their hosts quickly and the released progeny decay in the environment for the remainder of the activity period. Thus, progeny released early in the host activity period are more likely to die in the environment prior to contacting a naive host in the following season. An intermediate virulence level that allows parasites to kill their host prior to the end of the activity period, but not so quickly that the progeny produced are likely to decay in the environment, result in the greatest evolutionary fitness.

The optimal virulence level increases linearly with decreases in the duration of host activity (Figure 2). Virulent parasites in environments where host activity periods are short minimize the cost of not producing progeny from infected hosts and do not incur the costs of progeny decaying in the environment. By contrast, environments where host activity periods are long favor parasites with a long incubation period to limit the cost of progeny decay due to environmental exposure while still killing hosts prior to the end of the season. The optimal level of virulence in all environmental scenarios results in parasite-induced host death just prior to the end of the seasonal activity period. The linear increase in optimal virulence as season length decreases suggests that parasite fitness is optimized when host death occurs at a fixed time before the end of the season.

**Figure 2.**
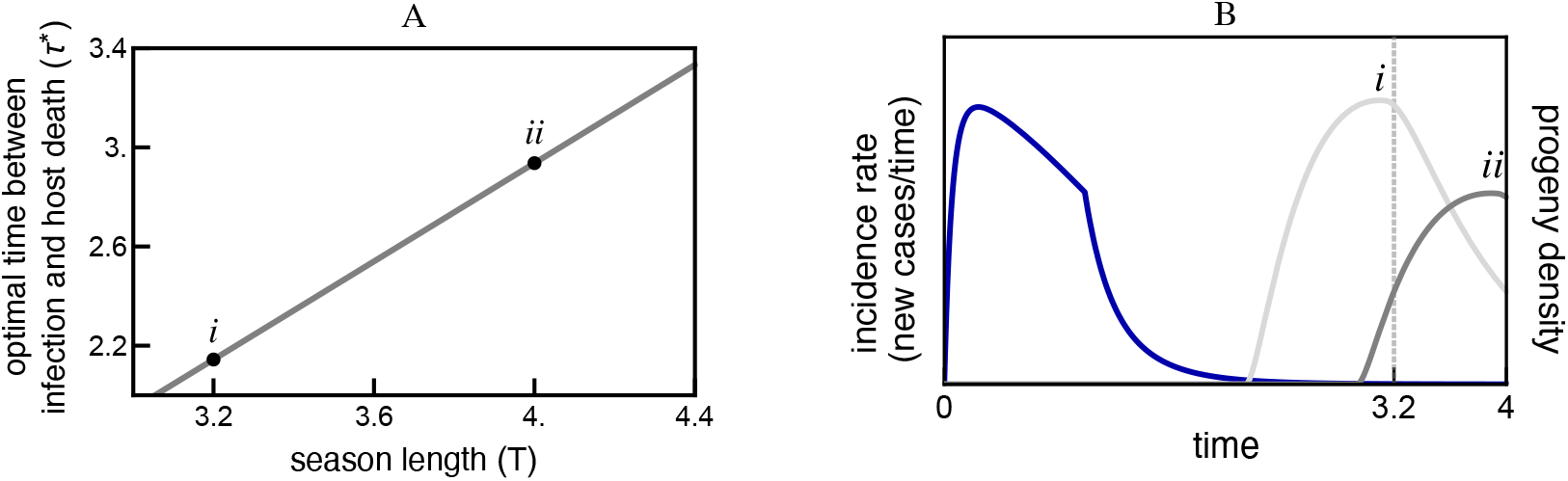
Host seasonality is sufficient to select an intermediate virulence strategy. A. The temporal duration between infection and host death (*τ*^∗^) always evolves to a value that is greater than 0 (extreme virulence) and less than the season length (extremely low virulence); the intermediate virulence strategy maximizes parasite fitness in environments where host activity is seasonal. The optimal parasite-induced host death rate results in host death and progeny release shortly before the end of the season (*t* = *T*). The release of progeny just prior to the end of the season limits the decay of progeny due to environmental exposure while avoiding progeny dying within their host at the end of the season. *i* and *ii* are representative examples of optimal virulence strategies in environments with shorter (*T* = 3.2) or longer (*T* = 4) host activity periods, respectively. *τ* ^∗^ is found using equation (4a) when there is no trade-off between transmission and virulence. **B**. Higher parasite virulence is favored in environments with limited host activity periods. Parasites with greater virulence produce more progeny that survive to the end of the season when seasons are short. That is, the density of the more virulent progeny (*i*) at *T* = 3.2 is greater than the density of the less virulent progeny (*ii*). The more virulent parasite kill their hosts quickly such that few infected hosts survive to the end of the season and the progeny released spend little time in the environment. By contrast, less virulent parasites (*ii*) often fail to kill their hosts and release progeny prior to the end of short activity periods (*T* = 3.2). Longer seasons (*T* = 4) favor less virulent parasites (*ii*) as they kill their hosts closer to the end of the season such that fewer of their released progeny decay in the environment (*ii*) than the progeny of the more virulent parasites that are released earlier in the season (*i*). The blue line represents the incidence rate of new infections; *t*_*l*_ = 1; all other parameters found in Table 1.

Variation in the time at which each host first becomes active during the activity period also impacts the virulence levels that maximize parasite fitness (Figure 3). Synchronous host emergence results in a rapid and early spike in infection incidence due to the simultaneous availability of susceptible hosts and the abundance of free parasites. The long duration between host infection and the end of the activity period favors low virulence parasites that kill their host near the end of the season (Figure 3A, *i*). Variability in the time at which each susceptible host initially becomes active decreases the average time between infection and the end of the season, thus favoring more virulent parasites (Figure 3A, *ii*). That is, the large proportion of infections that occur later in the season require higher virulence to be able to release progeny before the activity period ends. This higher virulence level comes at the cost of progeny from hosts infected early in the season decaying in the environment. Thus, the number of progeny that survive to the next season decreases with increasing variation in host emergence times (Figure 3B).

**Figure 3.**
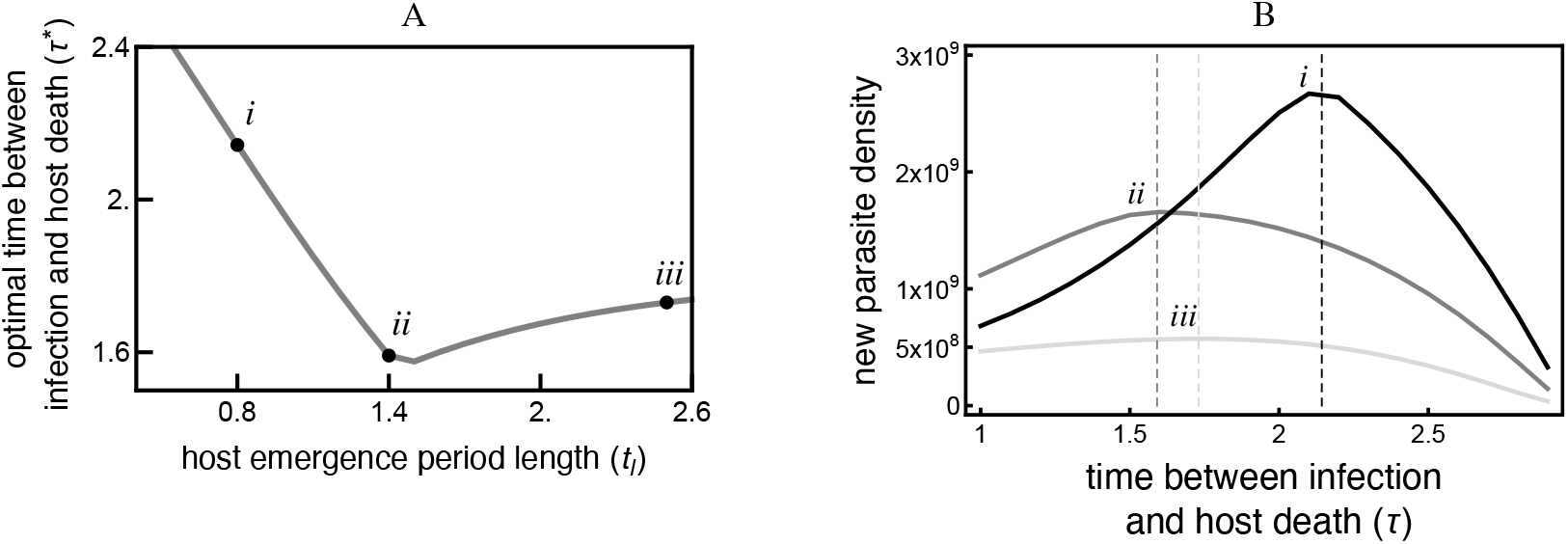
The variation in host emergence timing impacts the optimal virulence strategy. A. Parasites with lower virulence are favored in environments where nearly all hosts emerge simultaneously (*i*). Progeny from the low virulence parasites are released nearly simultaneously just prior to the end of the season. High virulence parasites are favored in environments where host emergence period length is moderate (*ii*). Moderate variation in host emergence decreases the average time between infection and the end of the season and favors parasites with a high virulence strategy such that few infected hosts survive to the end of the season. Parasites in environments where host emergence variation is very high maximize the number of progeny that survive to the next season by using a moderate virulence strategy (*iii*). Parasites in these environments suffer the costs of hosts that are infected later in the season not releasing progeny as well as progeny decay in the environment when released from early-infected hosts. A moderate virulence strategy allows hosts infected around the mid-season peak in incidence to release progeny while limiting the decay of these progeny. *τ* ^∗^ is found using equation (4a) when there is no trade-off between transmission and virulence. **B**. Equilibrium density of parasites with the optimal virulence strategy for their environment decreases with increasing variation in host emergence timing. Optimal virulence results in peak equilibrium in new parasites density, indicated by the vertical lines. *T* = 3; other parameters found in Table 1.

High variability in host emergence timing results in an optimal virulence strategy that is much greater than in environments with synchronous host emergence, but lower than in environments with a moderate distribution (Figure 3). That is, increasing variation in host emergence timing favors parasites with higher virulence, but only when variation in host emergence timing is moderate. In environments where the variation in host emergence timing is high, increasing variation in host emergence timing favors parasites with slightly lower virulence (Figure 3A, *iii*). Lower virulence is favored in high emergence variability environments because the number of new infections occurring late in the season, where high virulence would be advantageous, are relatively rare due to small parasite population sizes at the beginning of the season and parasite decay during the season. Initial parasite population sizes are smaller in environments with broadly distributed host emergence timing as fewer total hosts are infected because infection probability is density dependent, and thus fewer progeny are produced. Most parasites that find a susceptible host do so early in the season resulting in additional decreases to the already small parasite population size. The optimal virulence strategy allows parasites that infect hosts around the peak of new infections - occurring mid-season when susceptible host densities are greatest and parasite populations have not decayed substantially - to release progeny while limiting decay of these progeny. Parasites in environments where the distribution in host emergence times is very broad suffer the costs of both decay of the progeny released by early-infected hosts and the cost of late infected hosts not releasing progeny, collectively causing these environments to maintain low densities of moderately virulent pathogens (Figure 3B, *iii*).

Mechanistic virulence-transmission trade-offs can modify the optimal virulence strategy in seasonal environments but are not necessary for natural selection to favor intermediate virulence phenotypes. The optimal virulence strategy is slightly lower in models that include a trade-off where duration of infection is positively correlated with progeny production than in models with the same phenological parameters that do not include the trade-off (Figure 4). Including this trade-off increases the fitness benefit of longer-duration infections to a greater extent than the costs associated with infected host mortality not caused by the parasite. By contrast, the optimal virulence strategy is greater in models that include a trade-off where duration of infection is negatively correlated with progeny production than in similar models without the trade-off (Figure 4). Including this trade-off increases the fitness benefit of shorter-duration infections despite the added costs of greater parasite decay due to environmental exposure. Including mechanistic trade-offs modifies the selection pressures on virulence strategies but are not essential for an intermediate virulence strategy to be optimal in seasonal environments.

**Figure 4.**
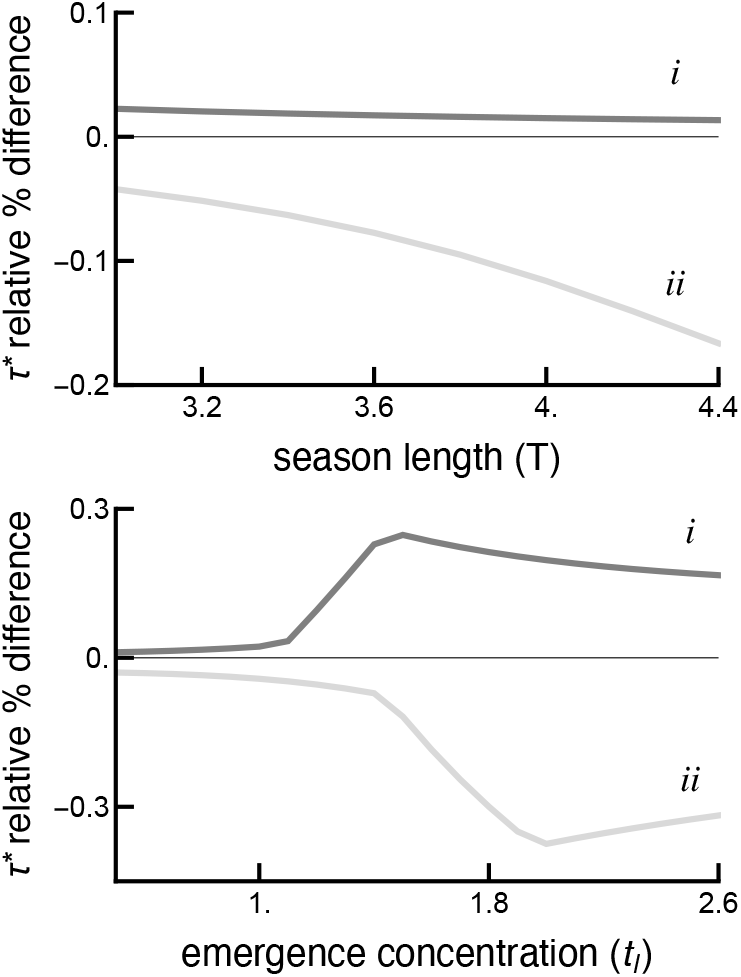
Mechanistic transmission-virulence trade-offs shift the optimal virulence strategy but are not necessary to favor intermediate virulence in environments with seasonal host activity. The optimal virulence level for parasites in which longer durations of infection result in *more* progeny is slightly lower than for parasites that are not constrained with this mechanistic trade-off in the same environment (*i*). This mechanistic trade-off elevates the fitness benefit of longer duration infections by compensating for the cost of infected hosts dying without releasing progeny. The optimal virulence level for parasites in which longer infection durations result in *fewer* progeny is greater than for parasites without this trade-off in the same seasonal environments (*ii*). This mechanistic trade-off elevates the fitness benefit of shorter duration infections despite the cost of greater progeny decay in the environment. *τ* ^∗^ was found using equation (4a) when there is no trade-off between transmission and virulence and then compared to *τ* ^∗^ constrained by a trade-off with transmission. Trade-off for *i* : *β*(*τ*) = 99(*τ* + 0.5), trade-off for *ii* : *β*(*τ*) = 99(*−τ* + 4). All other parameters found in Table 1.

## Discussion

Nearly all theory developed to explain parasite virulence evolution has utilized mechanistic trade-offs between virulence and other traits important to parasite fitness (Alizon et al., 2009; Cressler et al., 2016). The results of this study show that seasonal host activity, in the absence of an assumed negative correlation between virulence and transmission, can account for the evolution of intermediate virulence in some specific situations. Both aspects of phenology, the duration of the host activity period and host emergence synchronicity, impact the virulence strategy that maximizes the evolutionary fitness of parasites. Although mechanistic trade-offs between virulence and transmission can shift the optimal virulence level as predicted by prior theory, these trade-offs are not essential for intermediate virulence to evolve in this system. The current demonstration that an ecological context is sufficient to select for intermediate virulence broadens the scope of factors that can explain the diversity of parasite virulence strategies. Thus, the evolution of intermediate virulence in natural systems may be governed by a mechanistic trade-off or by ecological factors in some systems.

Seasonal host activity can select for intermediate virulence by generating conflicting costs for releasing progeny too early or too late in the season. Low virulence is maladaptive for parasites in this system as they do not kill their host before the end of the season and create no progeny. High virulence is also maladaptive as progeny released early are more likely to die due to environmental exposure. The conflicting costs of not releasing progeny before the end of the season and releasing progeny too early in the season selects for intermediate virulence levels. Optimal virulence results in parasite-induced host death and the release of progeny slightly before the end of the host activity period.

The result predicting adaptive evolution towards intermediate virulence stands in contrast to many prior theoretical investigations of obligate-killer parasites. Prior models of obligate-killer parasites predict ever-increasing virulence in the absence of mechanistic trade-offs (Caraco and Wang, 2008; Ebert and Weisser, 1997; Levin, 1983; Sasaki and Godfray, 1999). In simple obligate-killer models, killing infected hosts as quickly as possible is expected to maximize fitness as the early release of progeny permits infection of additional susceptible hosts resulting in a rapid exponential increase of parasites in the system. To date, only mechanistic trade-offs between virulence and transmission-associated factors as well as development time constraints have been demonstrated to constrain maximal virulence in obligate-killer parasite models (Ben-Ami, 2017; Caraco and Wang, 2008; Ebert and Weisser, 1997; Jensen et al., 2006; Wang, 2006). In contrast, our results indicate that host phenology can create conditions that favor intermediate virulence in obligate-killer parasites even if a negative correlation between virulence and transmission is not included in the model. In the current model, intermediate virulence is favored as seasonal host absence increases the evolutionary benefit of remaining within hosts in order to reduce deaths in the free-living stage caused by environmental exposure (Reece et al., 2017).

Variation in host emergence synchronicity impacts the optimal virulence strategy of parasites in this system. High parasite virulence is favored at low host emergence synchronicity. Low emergence synchronicity slows incidence by decreasing both the rate hosts emerge and parasite equilibrium density. When more infections occur later in the season, parasites have less time to release new parasites before the end of the season. High parasite virulence is adaptive because hosts have a low expected life span at the time of infection. This result is analogous to the prediction that high host mortality drives the evolution of high virulence (RM Anderson and May, 1982; Cooper et al., 2002; Gandon et al., 2001; Lenski and May, 1994). The timing of host activity can thus lead to the evolution of high virulence in a similar manner to how host demography impacts virulence.

The seasonal activity patterns of species with non-overlapping generations may have large impacts on the virulence strategies of the parasites they host. For example, parasites and parasitoids of univoltine insects that complete one round of infection per host generation may maximize their fitness by releasing progeny just prior to the end of the season (Baltensweiler et al., 1977; Bilimoria, 1991; Campbell, 1975; Delucchi, 1982; Dwyer, 1994; Dwyer and JS Elkinton, 1993; Kenis and Hilszczanski, 2007; Woods and J Elkinton, 1987). The theoretical expectations presented here can be tested empirically by measuring the virulence strategies of parasites across the natural diversity of phenological patterns observed over the geographical range of many insect species. Similarly, experiments could rigorously assess the impact of both season length and host emergence variability on the fitness of parasites with different levels of virulence.

The prediction that shorter host activity periods can drive greater virulence is comparable to how the virulence of different *Theileria parva* strains varies between regions. High within-host densities permit a virulent *T. parva* strain to be reliably transmitted to feeding nymphal tick vectors shortly after being infected by the adult stage in regions where the activity patterns of the two tick life stages overlap (Norval et al., 1991; Ochanda et al., 1996; Randolph, 1999). In contrast, the virulent strain is absent in regions where nymphal and adult activity is asynchronous while a less virulent strain that persists in hosts longer is maintained (Norval et al., 1991; Randolph, 1999). Thus, the prediction that the length of the host activity period is inversely correlated with virulence coincides with empirical observations of the distribution of *T. parva* strains.

Several features of the current model can be altered to investigate more complex impacts of phenology on virulence evolution. For example, relaxing the assumption of a constant host population size may result in a feedback between parasite fitness and host demography with consequences for population dynamics (Hilker et al., 2020). Additionally, parasite virulence evolution may select for alternative host phenological patterns that in turn select for parasite traits with lower impacts on host fitness. We will extend the current model to address these questions in future studies.

The model presented applies to obligate-killer parasites that complete one round of infection per season (monocyclic) in hosts that have non-overlapping generations. Currently, there is no evidence that disease systems that violate these assumptions can select for intermediate virulence without including a mechanistic trade-off. Nevertheless, several prior models that included both host seasonality and mechanistic trade-offs found qualitatively similar results as those presented here despite relaxing one or more of the strict assumptions in this model (Berg et al., 2011; King et al., 2009; Sorrell et al., 2009), suggesting that phenology can have a large impact on virulence outcomes. For example, longer seasons or longer periods between seasons have been shown to select for lower virulence in polycyclic parasites in seasonal environments (Berg et al., 2011; Sorrell et al., 2009), similar to the results presented here. Similarly, explicitly modeling parasite growth rates within hosts, which underlie the correlation between virulence and instantaneous transmission rates, selects for intermediate virulence levels that maximize transmission rates during host activity periods (King et al., 2009). By contrast, assuming that virulence levels are mechanistically associated with host density results in selection for higher virulence in seasonal environments (Donnelly et al., 2013). Future studies incorporating one or more of these competing forces with environmental decay of progeny could be sufficient to select for intermediate virulence in the absence of an assumed mechanistic trade-off.

Some of the strict model assumptions can likely be relaxed without altering the result that phenology can be sufficient to select for intermediate virulence strategies. Relaxing the obligate-killer assumption may result in the same qualitative result that intermediate virulence is adaptive in some cases. For example, longer latency periods that result in progeny release near the end of the season would still be adaptive for parasites that reduce host fecundity or increase host death rate, even if there is no correlation between the virulence level and instantaneous or life-time transmission. Longer latency periods are equivalent to lower virulence in this type of system as infected hosts have more time to reproduce and thus higher fitness. This extension is not expected to qualitatively alter the results if the parasite transmission period is short relative to the season length. Many parasite-host systems conform to the assumptions of this model extension such as monocyclic plant pathogens (*e*.*g*. soil-borne plant pathogens, demicyclic rusts, post-harvest diseases), and many diseases systems infecting univoltine insects (Crowell, 1934; Gaulin et al., 2007; Hamelin et al., 2011; Holuša and Lukášová, 2017; Zehr et al., 1982).

The importance of parasite virulence to both host-parasite interactions and public health policy has resulted in a concentrated research effort on virulence evolution. Nearly all theoretical research to date has incorporated a mechanistic trade-off between virulence and transmission rates or infection duration, a hypothesis which is still essential to explain the evolution of intermediate virulence in most disease systems. However, ecological factors such as seasonal host activity or spatial structuring provide alternative theoretical frameworks that may account for virulence strategies in some natural systems (Boots and Sasaki, 1999; Kerr et al., 2006). Future work that identifies and empirically validates ecological factors that influence virulence evolution would be useful for predicting outbreaks of highly virulent parasites.

## Code and data availability

Code is available on the Github repository: https://github.com/hanneloremac/Host-phenology-drives-the-evolution-of-intermediate-parasite-virulence

## Acknowledgements

Version 8 of this preprint has been peer-reviewed and recommended by *Peer Community In Evolutionary Biology* (https://doi.org/10.24072/pci.evolbiol.100129)

## Conflict of interest disclosure

The authors of this preprint declare that they have no financial conflict of interest with the content of this article. Dustin Brisson and Erol Akçay are recommenders for PCI Evolutionary Biology

## Appendix A

In Appendix A we find analytical solutions for equations (1a-c) from the main text to study parasite fitness given different host phenological patterns.

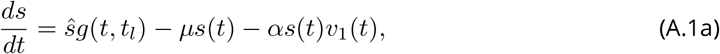

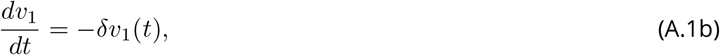

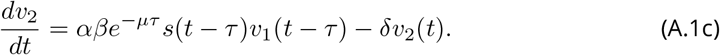

with initial conditions: *s*(0) = 0, *v*_1_(0^+^) = *v*_2_(0^*−*^), *v*_2_(*τ*) = 0.

(A.1a-c) is solved analytically by describing host emergence using a uniform distribution

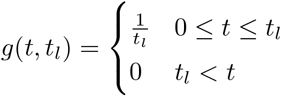

To solve the dynamics during the host*’*s activity period, we first find the analytical solution for *v*_1_(*t*):

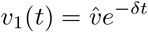

We then use *v*_1_(*t*) to find the time-dependent solution for *s*(*t*). We can then plug the time-dependent solution for *s*(*t*) to find the time-dependent solution for *v*_2_(*t*). Only parasites that infect hosts from 0 *< t < T − τ* have enough time to kill hosts and release progeny before the end of the season. For *τ < T − t*_*l*_, parasites that infect hosts during host emergence (0 *< t* ≤ *t*_*l*_) have time to kill hosts and release progeny before the end of the season as well as some parasites who infect hosts after host emergence has ended (*t > t*_*l*_). For *τ > T − t*_*l*_, only some parasites that infect hosts during host emergence (0 *< t* ≤ *t*_*l*_) have time to kill hosts and release progeny before the end of the season. Thus, two separate solutions are required depending on whether *τ* is greater or less than *T − t*_*l*_. We first consider the case where *τ < T − t*_*l*_:

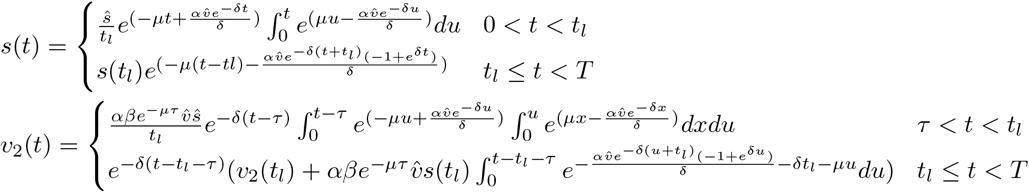

where *s*(*t*_*l*_) and *v*_2_(*t*_*l*_) are the densities of *s* and *v*_2_ when the emergence period of *s* ends.

For *τ > T − t*_*l*_, only some of the parasites that infect hosts from 0 *< t < t*_*l*_ have enough time to kill hosts and release progeny before the end of the season. *v*_2_(*t*) are thus only produced from infections that occurred from 0 *< t < t*_*l*_. The solution for *v*_2_(*t*) in this case is

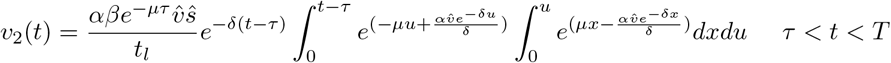

A parasite introduced into a naive host population persists or goes extinct depending on the host emergence period length and season length. The stability of the parasite-free equilibrium when *τ < T − t*_*l*_ is determined by the production of *v*_2_ resulting from infection of *s* given by

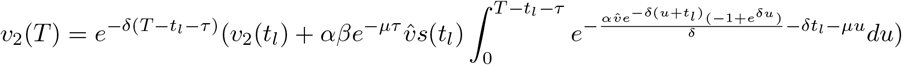

When *τ > T − t*_*l*_, the stability of the parasite-free equilibrium is determined by

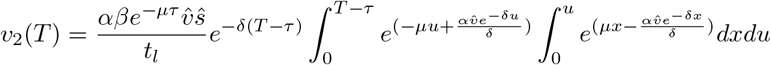

The parasite-free equilibrium is unstable and the parasite will persist in the system if the density of *v*_2_ produced by time *T* is greater than or equal to 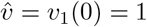 introduced at the beginning of the activity period of *s* (*i*.*e. v*_2_(*T*) ≥ 1, modulus is greater than unity). This expression is a measure of a parasite*’*s fitness when rare for different host phenological patterns given *τ > T − t*_*l*_.

## Appendix B

In Appendix B we find analytical solutions for equations 2a-e from the main text to study the evolution of parasite virulence given different host phenological patterns.

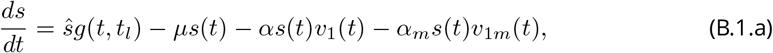

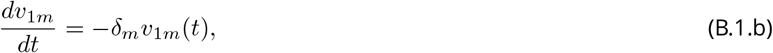

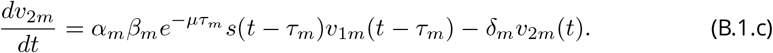

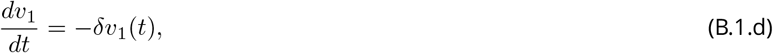

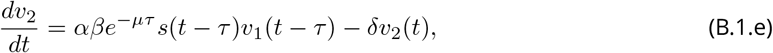

with initial conditions: *s*(0) = 0, *v*_1*m*_(0^+^) = *v*_2*m*_(0^*−*^), *v*_2*m*_(*τ*) = 0, *v*_1_(0^+^) = *v*_2_(0^*−*^), *v*_2_(*τ*) = 0. *m* subscripts refer to the invading mutant parasite and its corresponding traits.

Again, separate solutions for (B.1.a-c) are required depending on whether *τ* and *τ*_*m*_ are greater or less than *T − t*_*l*_. The length of *τ* relative to *T* and *t*_*l*_ determines 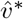 while the length of *τ*_*m*_ relative to *T* and *t*_*l*_ determines the within-season dynamics of the mutant parasite. The solutions to all cases can be found in the code on Github (see *“*Code and data availability*”* in the main text for the link). We first show the solution to the case when *τ*_*m*_ *< T − t*_*l*_:

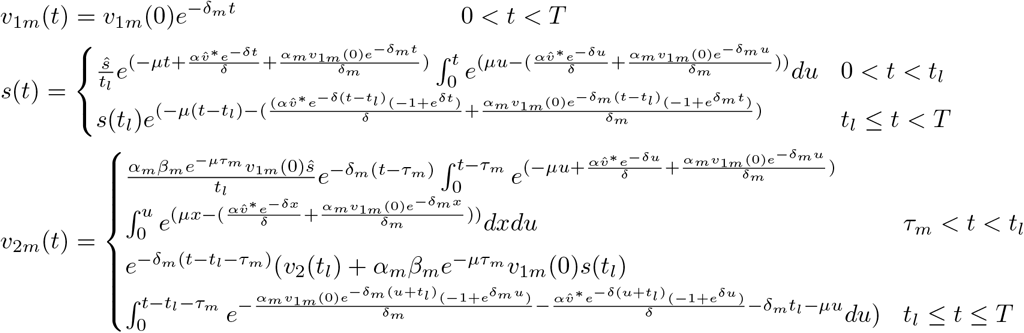

When *τ*_*m*_ *> T − t*_*l*_, the solution for *v*_2*m*_(*t*) is

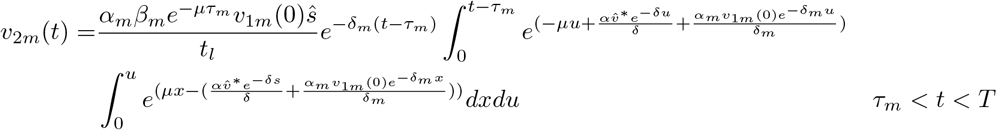

The invasion fitness of a rare mutant parasite is given by the density of *v*_2*m*_ produced by the end of the season. When *τ*_*m*_ *< T − t*_*l*_, the mutant parasite invades in a given host phenological scenario if the density of *v*_2*m*_ produced by time *T* is greater than or equal to the initial *v*_1*m*_(0) = 1 introduced at the start of the season (*v*_2*m*_(*T*) ≥ 1), following

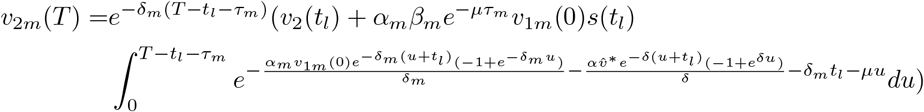

When *τ*_*m*_ *> T − t*_*l*_, the mutant parasite invades in a given host phenological scenario if the density of *v*_2*m*_ produced by time *T* is greater than or equal to the initial *v*_1*m*_(0) = 1 introduced at the start of the season (*v*_2*m*_(*T*) ≥ 1), following

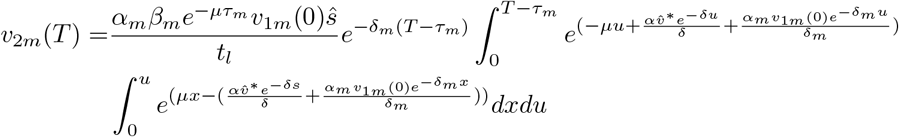

We use *v*_2*m*_(*T*) to find optimal virulence for a given host phenological scenario by finding the trait value that maximizes *v*_2*m*_(*T*). That is, the virulence trait, *τ* ^∗^, that satisfies

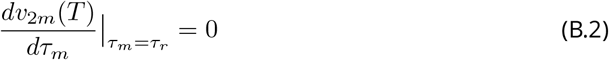

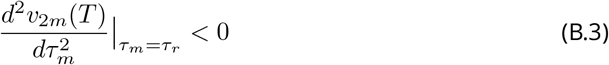

For all phenological patterns, we found that *τ* ^∗^ is uninvadable *i*.*e*. condition (B.3) is satisfied.

For certain phenological patterns, *τ*_*m*_ switches from *T − t*_*l*_ *< τ*_*m*_ to *T − t*_*l*_ *> τ*_*m*_ as it evolves. When *t*_*l*_ is small, optimal virulence, *τ* ^∗^, is short relative to *T* and *t*_*l*_. Thus when *t*_*l*_ is short, *τ* ^∗^ times parasite-induced death to begin *after* all hosts have finished emerging (*i*.*e*. the solution where *T − t*_*l*_ *> τ*_*m*_ is required to find *τ* ^∗^). For large *t*_*l*_, the value of *τ* ^∗^ that optimizes parasite fitness initiates parasite-induced host death *before* all hosts have finished emerging (*i*.*e*. the solution where *T − t*_*l*_ *< τ*_*m*_ is required to find *τ* ^∗^). We found the value of *t*_*l*_ that requires a switch from the solution for *T − t*_*l*_ *> τ*_*m*_ to the solution for *T − t*_*l*_ *< τ*_*m*_ to find *τ* ^∗^ numerically using Mathematica. We switched which solution we used to find *τ* ^∗^ when the value of *τ* ^∗^ that satisfied (B.2) no longer met the inequality. For example, for long *t*_*l*_, the solution for *T − t*_*l*_ *> τ*_*m*_ returned *T − t*_*l*_ *< τ* ^∗^. When this occurred we switched to using the solution for *T − t*_*l*_ *< τ*_*m*_ to find *τ* ^∗^.

To study the impact of mechanistic trade-offs between transmission and virulence on virulence evolution, we assume that the number of parasites produced at host death is a function of the time between infection and host death (*β*(*τ*)). This is done in Figures 3A and 3B where a mechanistic trade-off is assumed to exist between *τ* and *β* in (B.2). The same approach as described above is used to determine which solution correctly specifies *τ* ^∗^.

## Notes

### Competing Interest Statement

The authors have declared no competing interest.

### Summary of Updates

Version 8 of this preprint has been peer-reviewed and recommended by Peer Community In Evolutionary Biology (https://doi.org/10.24072/pci.evolbiol.100129)

## References

Alizon S, A Hurford, N Mideo, and M Van Baalen (2009). Virulence evolution and the trade-off hypothesis: history, current state of affairs and the future. J Evol Biol 22, 245–259.

Altizer S, A Dobson, P Hosseini, P Hudson, M Pascual, and P Rohani (2006). Seasonality and the dynamics of infectious diseases. Ecology Letters 9, 467–484.

Anderson J, D Inouye, A McKinney, R Colautti, and T Mitchell-Olds (2012). Phenotypic plasticity and adaptive evolution contribute to advancing flowering phenology in response to climate change. Proc Biol Sci 279, 3843–3852.

Anderson RM and RM May (1982). Coevolution of hosts and parasites. Parasitology 85, 411– 426.

Anderson RM and RM May (1981). The population dynamics of microparasites and their invertebrate hosts. Philosophical Transactions of the Royal Society of London. B, Biological Sciences 291, 451–524.

Baltensweiler W, G Benz, P Bovey, and V Delucchi (1977). Dynamics of larch bud moth populations. Annual Review of Entomology 22, 79–100.

Ben-Ami F (2017). The virulence–transmission relationship in an obligate killer holds under diverse epidemiological and ecological conditions, but where is the tradeoff? Ecology and evolution 7, 11157–11166.

Berg F van den, N Bacaer, JA Metz, C Lannou, and F van den Bosch (2011). Periodic host absence can select for higher or lower parasite transmission rates. Evolutionary Ecology 25, 121–137.

Biere A and SJ Honders (1996). Impact of Flowering Phenology of Silene alba and S. dioica on Susceptibility to Fungal Infection and Seed Predation. Oikos 77, 467–480.

Bilimoria SL (1991). The biology of nuclear polyhedrosis viruses. Viruses of invertebrates, 1–72.

Boots M and A Sasaki (1999). ‘Small worlds’ and the evolution of virulence: infection occurs locally and at a distance. Proceedings of the Royal Society of London. Series B: Biological Sciences 266, 1933–1938.

Campbell RW (1975). The gypsy moth and its natural enemies. 381. US Department of Agriculture, Forest Service.

Caraco T and IN Wang (2008). Free-living pathogens: life-history constraints and strain competition. Journal of theoretical biology 250, 569–579.

Cooper V, M Reiskind, J Miller, K Shelton, B Walther, J Elkinton, and P Ewald (2002). Timing of transmission and the evolution of virulence of an insect virus. Proc Biol Sci 269, 1161–1165.

Cressler CE, DV McLEOD, C Rozins, J van den Hoogen, and T Day (2016). The adaptive evolution of virulence: a review of theoretical predictions and empirical tests. Parasitology 143, 915– 930.

Crowell IH (1934). The hosts, life history and control of the cedar-apple rust fungus Gymnosporangium juniperi-virginianae Schw. Journal of the Arnold Arboretum 15, 163–232.

Delucchi V (1982). Parasitoids and hyperparasitoids of Zeiraphera diniana [Lep., Tortricidae] and their pole in population control in outbreak areas. Entomophaga 27, 77–92.

Donnelly R, A Best, A White, and M Boots (2013). Seasonality selects for more acutely virulent parasites when virulence is density dependent. Proceedings of the Royal Society B: Biological Sciences 280, 20122464.

Dwyer G (1994). Density dependence and spatial structure in the dynamics of insect pathogens. The American Naturalist 143, 533–562.

Dwyer G and JS Elkinton (1993). Using simple models to predict virus epizootics in gypsy moth populations. Journal of Animal Ecology, 1–11.

Ebert D and WW Weisser (1997). Optimal killing for obligate killers: the evolution of life histories and virulence of semelparous parasites. Proceedings of the Royal Society of London. Series B: Biological Sciences 264, 985–991.

Elzinga JA and e al et (2007). Time after time: flowering phenology and biotic interactions. Trends in Ecology and Evolution.

Forrest J and AJ Miller-Rushing (2010). Toward a synthetic understanding of the role of phenology in ecology and evolution. Philosophical transactions of the Royal Society of London. Series B, Biological sciences 365, 3101–3112.

Gandon S, VA Jansen, and M Van Baalen (2001). Host life history and the evolution of parasite virulence. Evolution 55, 1056–1062.

Gaulin E, C Jacquet, A Bottin, and B Dumas (2007). Root rot disease of legumes caused by Aphanomyces euteiches. Molecular Plant Pathology 8, 539–548.

Geritz SA, G Mesze, JA Metz, et al. (1998). Evolutionarily singular strategies and the adaptive growth and branching of the evolutionary tree. Evolutionary ecology 12, 35–57.

Gethings O, H Rose, S Mitchell, J Van Dijk, and E Morgan (2015). Asynchrony in host and parasite phenology may decrease disease risk in livestock under climate warming: Nematodirus battus in lambs as a case study. Parasitology 142, 1306–1317.

Hamelin FM, M Castel, S Poggi, D Andrivon, and L Mailleret (2011). Seasonality and the evolutionary divergence of plant parasites. Ecology 92, 2159–2166.

Hamer SA, GJ Hickling, JL Sidge, ED Walker, and JI Tsao (2012). Synchronous phenology of juvenile Ixodes scapularis, vertebrate host relationships, and associated patterns of Borrelia burgdorferi ribotypes in the midwestern United States. Ticks and Tick-borne Diseases 3, 65–74.

Hamilton W (1964a). The genetical evolution of social behaviour. I. Journal of theoretical biology. 7. ISSN: 0022-5193.

Hamilton W (1964b). The genetical evolution of social behaviour. II. Journal of theoretical biology. 7. ISSN: 0022-5193.

Hilker F, T Sun, L Allen, and F Hamelin (2020). Separate seasons of infection and reproduction can lead to multi-year population cycles. Journal of theoretical biology 489, 110158.

Holuša J and K Lukášová (2017). Pathogen’s level and parasitism rate in Ips typographus at high population densities: importance of time. Journal of Applied Entomology 141, 768–779.

Jensen KH, T Little, A Skorping, and D Ebert (2006). Empirical support for optimal virulence in a castrating parasite. PLoS Biol 4, e197.

Kenis M and J Hilszczanski (2007). Natural enemies of Cerambycidae and Buprestidae infesting living trees. In: Bark and wood boring insects in living trees in Europe, a synthesis. Springer, pp. 475–498.

Kerr B, C Neuhauser, BJM Bohannan, and AM Dean (2006). Local migration promotes competitive restraint in a host–pathogen ‘tragedy of the commons’. Nature 442, 75–78.

King AA, S Shrestha, ET Harvill, and ON Bjørnstad (2009). Evolution of acute infections and the invasion-persistence trade-off. The American Naturalist 173, 446–455.

Lenski RE and RM May (1994). The Evolution of Virulence in Parasites and Pathogens: Reconciliation Between Two Competing Hypootheses. Journal of Theoretical Biology 169, 253– 265.

Levin B (1983). Coevolution in bacteria and their viruses and plasmids. Coevolution, 99–127.

Lion S and JA Metz (2018). Beyond R0 maximisation: on pathogen evolution and environmental dimensions. Trends in ecology & evolution 33, 458–473.

Lustenhouwer N, R Wilschut, J Williams, W van der Putten, and J Levine (2018). Rapid evolution of phenology during range expansion with recent climate change. Glob Chang Biol 24, e534–e544.

MacDonald H, E Akçay, and D Brisson (2020). The role of host phenology for parasite transmission. Theoretical Ecology.

Martinez ME (2018). The calendar of epidemics: Seasonal cycles of infectious diseases. PLOS Pathogens 14, e1007327–15.

McDevitt-Galles T, WE Moss, DM Calhoun, and PT Johnson (2020). Phenological synchrony shapes pathology in host–parasite systems. Proceedings of the Royal Society B 287, 20192597.

Metz JA, RM Nisbet, and SA Geritz (1992). How should we define ‘fitness’ for general ecological scenarios? Trends in ecology & evolution 7, 198–202.

Norval R, J Lawrence, A Young, BD Perry, T Dolan, and J Scott (1991). Theileria parva: influence of vector, parasite and host relationships on the epidemiology of theileriosis in southern Africa. Parasitology 102, 347–356.

Novy A, S Flory, and J Hartman (2013). Evidence for rapid evolution of phenology in an invasive grass. J Evol Biol 26, 443–450.

Ochanda H, A Young, C Wells, G Medley, and BD Perry (1996). Comparison of the transmission of Theileria parva between different instars of Rhipicephalus appendiculatus. Parasitology 113, 243–253.

Ogden NH, G Pang, HS Ginsberg, GJ Hickling, RL Burke, L Beati, and JI Tsao (2018). Evidence for Geographic Variation in Life-Cycle Processes Affecting Phenology of the Lyme Disease Vector Ixodes scapularis (Acari: Ixodidae) in the United States. Journal of Medical Entomology 55, 1386–1401.

Park J (2019). Cyclical environments drive variation in life-history strategies: a general theory of cyclical phenology. Proc Biol Sci 286, 20190214.

Pau S, EM Wolkovich, BI Cook, TJ Davies, NJB Kraft, K Bolmgren, JL Betancourt, and EE Cleland (2011). Predicting phenology by integrating ecology, evolution and climate science. Global Change Biology 17, 3633–3643.

Randolph SE (1999). Tick ecology: processes and patterns behind the epidemiological risk posed by ixodid ticks as vectors. Parasitology 129, S37–S65.

Reece SE, KF Prior, and N Mideo (2017). The life and times of parasites: rhythms in strategies for within-host survival and between-host transmission. Journal of biological rhythms 32, 516–533.

Sasaki A and H Godfray (1999). A model for the coevolution of resistance and virulence in coupled host–parasitoid interactions. Proceedings of the Royal Society of London. Series B: Biological Sciences 266, 455–463.

Smith T (1904). Some problems in the life history of pathogenic microorganisms. Science 20, 817–832.

Sorrell I, A White, AB Pedersen, RS Hails, and M Boots (2009). The evolution of covert, silent infection as a parasite strategy. Proceedings of the Royal Society B: Biological Sciences 276, 2217–2226.

Wang IN (2006). Lysis timing and bacteriophage fitness. Genetics 172, 17–26.

White MT, G Shirreff, S Karl, AC Ghani, and I Mueller (Mar. 2016). Variation in relapse frequency and the transmission potential of Plasmodium vivax malaria. Proceedings of the Royal Society B: Biological Sciences 283, 20160048–9.

White NJ (2011). Determinants of relapse periodicity in Plasmodium vivax malaria. Malaria journal 10, 1–36.

Woods S and J Elkinton (1987). Biomodal patterns of mortality from nuclear polyhedrosis virus in gypsy moth (Lymantria dispar) populations. Journal of Invertebrate Pathology 50, 151–157.

Zehr EI et al. (1982). Control of brown rot in peach orchards. Plant disease 66, 1101–1105.

